# Accounting for allelic diversity and multicopy gene detection improves the accuracy of antibiotic resistance genotypic determination

**DOI:** 10.64898/2026.06.08.729070

**Authors:** Neris García-Gonzalez, Roser Ferragud, Beth Blane, Jee In Kim, M Estée Török, Ewan M Harrison, Theodore Gouliouris, Francesc Coll

## Abstract

**Background:** Genomic prediction of antimicrobial resistance (AMR) relies on the accurate detection of resistance genes or allelic variants of core genes from raw or assembled genomes sequences. For several bacterial species and antibiotics, AMR genotype-phenotype discrepancies are common, indicating that important sources of error remain unresolved. For *Enterococcus faecium*, we focused on identifying the sources of discrepancies for tetracycline resistance, for which genotypic detection had shown particularly low accuracy. We investigated the effect of structural variation in antibiotic resistance genes (ARGs)—including gene duplications, truncations, interruptions, and mixed configurations of complete and partial gene copies— as a source of genotype-phenotype discrepancies from short□read data. We conduct further extended investigations to other antibiotic families and into another bacterial species: *Escherichia coli*.

**Methods:** We analyzed collections of *E. faecium* and *E. coli* genomes, integrating high□quality complete assemblies, simulated Illumina short reads, and matched AMR phenotypic data. The integrity, copy number, and allelic diversity of ARGs were examined for multiple antibiotic classes, and their impact on ARG detection and accuracy of AMR determination was assessed using several commonly used bioinformatic tools (SRST2, ARIBA and AMRFinderPlus).

**Results:** For *E. faecium*, after ruling out the effect of specific *tet* allelic variants on tetracycline susceptibility, we found that the integrity and copy number of *tet*(M) had a major effect on detection accuracy. Duplicated and incomplete ARGs are also common in *E. faecium* genomes, particularly for macrolides (*erm*(B)) and aminoglycosides (*ant(6)-Ia* and *aph(3’)-IIIa*). In *E. coli*, similar patterns were observed for *tet*(A)*, erm*(B) and aminoglycoside□associated genes (*aph(3□)-IIIa* and *ant(6)-Ia*). Across ARGs in both species, short-read mapping methods wrongly reported interrupted genes as complete in some instances, while assembly□based methods often failed to resolve complete copies of duplicated genes. Detection accuracy improved when tools were adapted to account for gene integrity and when extended AMR databases incorporating species□specific alleles were included.

**Conclusions:** Our findings reveal that bioinformatic limitations in dealing with ARG copy number and completeness, and in accounting for allelic variation, underly a substantial source of genotype-phenotype errors, highlighting the need for improved AMR databases and bioinformatic tools that consider these factors to achieve reliable genomic prediction of AMR.

## Background

Accurate determination of antimicrobial resistance (AMR) phenotypes from genomic data is a cornerstone of modern AMR surveillance and diagnostics. Genotype-based approaches rely on the identification of known AMR determinants from raw sequence data or genome assemblies, assuming that the presence of an AMR genetic marker reliably indicates phenotypic resistance [1]. However, despite well-curated AMR databases and maintained bioinformatic pipelines, genotype-phenotype discrepancies remain common for many bacterial species and drug classes [2–6], limiting the routine implementation of genomic AMR genotypic inference.

Previous studies have revealed multiple explanations for genotype-phenotype discordance, including incomplete knowledge of the genetic basis of phenotypic resistance, particularly for newly developed drugs; variable phenotypic effects of different allelic variants in the same AMR genes (ARGs) [7]; silenced acquired ARGs [8]; and the influence of the bioinformatic pipeline and AMR databased used [9,10]. However, these factors do not fully account for all mismatches observed across large genomic datasets.

In *Enterococcus faecium*, a gut commensal and important nosocomial pathogen, resistance to most antibiotics can be accurately detected genotypically by current AMR databases and tools, but genotypic inference of resistance to tetracyclines, macrolides, and aminoglycosides phenotypes show low specificity (∼60–90%) [3]. Tetracycline resistance in *E. faecium* is primarily mediated by ribosomal protection proteins encoded by *tet(M)* and efflux pumps encoded by *tet*(L). *tet*(M) is among the most prevalent tetracycline resistance genes in this species and is often carried on mobile genetic elements (MGEs), which can undergo duplications, rearrangements, partial deletions, or interruptions that affect gene integrity [11].

Here, we hypothesized that structural variation in ARGs (completeness and duplication) could contribute to AMR genotype-phenotype discrepancies from *E. faecium* short□read data. We also evaluated how AMR database representativeness, in terms of ARG allele representation, affects the performance of genotypic inference. We then extended the analysis to other antibiotic classes and *E. coli* and showed that these factors contribute to inaccuracies in gene detection and AMR determination more broadly, i.e. not exclusive to *E. faecium* and tetracycline resistance. Our findings highlight that accounting for gene integrity, gene copy-number, and representativeness of ARG alleles can improve the accuracy of genomic ARG detection and, consequently, genotype-based phenotype inference.

## Methods

### Genome collection

#### E. faecium short-read dataset with phenotypic AST data

We analyzed a collection of 4,764 *E. faecium* isolates with previously curated Antimicrobial Susceptibility Testing (AST) and Whole Genome Sequencing (WGS) data [3] (Additional file 2: Table S1). Guidelines and Minimum Inhibitory Concentration (MIC) cutoffs used to interpret AST detailed in Table S2 in additional file 2. For the analysis of tetracycline resistance, we retained only Clade A isolates with available tetracycline AST data, yielding 2,610 genomes. Illumina raw reads were downloaded from the Sequence Read Archive (SRA) using *fastq-dump* from sra-toolkit *v2.9.6* [12], and corresponding assemblies were obtained from AllTheBacteria v2 [13]. Three isolates carrying partial *tet(M)* genes were resequenced using Oxford Nanopore (ONT) long-read sequencing to characterize their genomic contexts.

Genomic DNA was extracted using a QIAamp Mini DNA kit (QIAgen, Germany). Long-read sequencing was performed at the Wellcome Sanger Institute (WSI) using the ONT PromethION with R10.4.1 flow cells, and libraries were prepared using the Rapid Barcoding Kit SQK-RBK114.96. Basecalling was performed using Dorado v7.2.13 [14] in super-accuracy (SUP) simplex mode, followed by barcode-based demultiplexing. Initial quality control of the resulting FASTQ reads was performed using Kraken2 (v2.1.2) [15] with the standard database (k2_standard_20250402) to screen for potential taxonomic contamination prior to downstream assembly.

#### Hybrid assemblies

ONT long reads were combined with the existing short-read data using a Hybracter-inspired WSI internal workflow, a long-read first bacterial genome assembly pipeline [16]. Briefly, Hybracter-like workflow performs read filtering and subsampling using Filtlong v0.2.1[17], long-read assembly with Flye v2.9.4 [18], followed by polishing with Medaka v1.11.3 [19], reorientation of circular contigs using Dnaapler v0.8.0 [20], and plasmid assembly using Plassembler v1.6.2 [21]. With the long-read polished draft, short-read polishing is completed with Polypolish v0.6.0 [22], and Pypolca v0.3.1 [23,24], providing a final assembly in FASTA format. Assemblies are available under BioProject Accession PRJNA1456658.

#### E. faecium closed genomes dataset

A dataset of 897 complete (i.e. closed) *E. faecium* genomes was downloaded from the National Center for Biotechnology Information (NCBI) (July 2025). Assembly quality was assessed with CheckM v1.1.3 [25] and BUSCO v5.8.3 [26] using the “enterococcus_odb12” lineage dataset□. After stringent quality filtering, 823 genomes were retained for downstream analyses (Additional file 2: Table S3, Additional file 1: Figure S1). Assemblies were excluded if they did not meet the following ChecKM metrics: an N50 value of at least 2.4 Mb, no more than 12 contigs, contamination not exceeding 0.7%, and completeness of at least 99%. Additionally, assembly quality was further assessed using BUSCO markers, which are a set of core single□copy orthologs for a given lineage, in this case *Enterococcus*. Genomes were required to have at least 950 complete BUSCO markers, and 950 single-copy BUSCOs, while those with more than 7 duplicated BUSCOs, 8 fragmented BUSCOs, or 30 missing BUSCOs were removed.

### Short-read simulations and benchmarking of AMR detection tools

To test how gene completeness and duplications may influence ARG detection, we simulated perfect Illumina paired-end reads from closed genomes using *wgsim v1.20* [27] (6 million pairs, 150 bp reads, insert size 350 ± 50 bp, no sequencing errors) (Figure 1 A). Simulated reads were analyzed using four ARG detection tools: SRST2 v0.2.0 [28], ARIBA v2.14.6 [29], ResFinder [30], and AMRFinderPlus v3.12.8 [31], all with the same underlying ARG catalogue, the AMRFinderPlus database (version 2025-03-25.1), except for ResFinder which uses its own database. AMRFinderPlus was run on *de novo* assemblies generated with Shovill v1.1.0 [32]. The minimum coverage for SRST2 and AMRFinderPlus was reduced to 20% to enable detection of partial ARGs.

**Figure 1.**
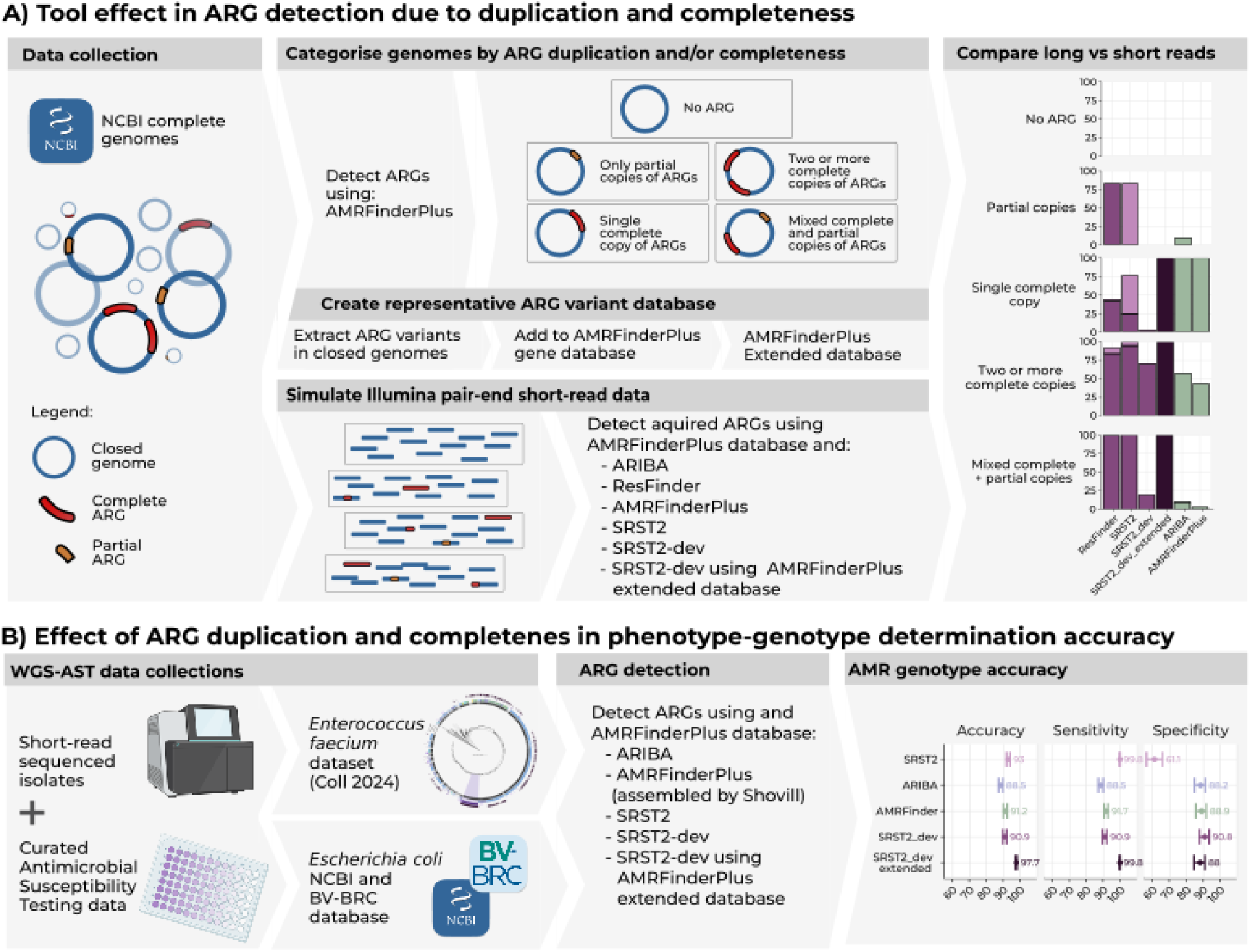
Methodological workflow for evaluating AMR gene detection accuracy and genotype-phenotype prediction. (A) Benchmarking of ARG detection from short□read data. Illumina□like short reads were simulated from a curated collection of closed *E. faecium* genomes from the NCBI. These reads were processed using four widely used ARG detection tools (SRST2, ARIBA, ResFinder, and AMRFinderPlus). The resulting ARG calls were compared directly against the ARG annotations of the same closed genomes to quantify each tool’s ability to correctly identify complete, partial, duplicated, or structurally interrupted ARGs. (B) GenotypeObased phenotype prediction for each tool was evaluated using two independent short□read clinical collections: *E. faecium* (n□=□4,507 genomes) and *E. coli* (n□=□7,437 genomes), each with AST results. ARG detection was compared to AST. Metrics including sensitivity, specificity, and overall diagnostic accuracy were calculated to quantify how structural variation (duplication, truncation, and allele diversity) in ARGs influences phenotype prediction in both species.

### SRST2 edition and AMR database extension

We developed a modified version of SRST2 (SRST2_dev) that preserves the original detection and allele scoring methods, but we added an additional “Truncated” flag, applied when more than 90% of reads at a given position were soft-clipped or when more than 75% were soft-clipped followed by a ≥3× drop in local coverage. The modified tool is available at https://github.com/NerisGarcia/srst2_NGG. In addition, we constructed extended ARG databases by supplementing the AMRFinderPlus database with all unique allelic variants identified in the closed-genome collection. For *E. faecium* we included allelic variants of *tet*(M), *erm*(B), *aph(30)*, and *ant(6)* whereas for *E. coli* we included variants for *tet*(A), *aph(6)*, *aph(3)*, *floR*, and *aadA2*.

### Assessing the diagnostic accuracy of genotypic AMR determinations

For each AMR detection tool tested, we evaluated the accuracy of genotypic inferred phenotypes using the AST phenotypic results as reference (Figure 1 B). We calculated commonly used metrics of diagnostic performance using the epi.tests function from the epiR v2.0.85 R package, including the following: diagnostic accuracy, sensitivity, and specificity. 95% confidence intervals were calculated using Wilson’s score.

### E. coli genome and AST datasets

We downloaded all complete closed *E. coli* genomes in the NCBI (by 21 July 2025). A stringent quality control process was applied to all genomes prior to downstream analyses using ChecKM and following qualibact suggested thresholds [33] (Additional file 2: Table S4). Assemblies were included if they had an N50 value of at least 20,000 bp, no more than 700 contigs, contamination not exceeding 16%, completeness of at least 95%, and GC content between 50.0% to 52.0%, a total coding sequences between 3900 and 6500 and a genome size between 4.1 to 6.3 Mb (Additional file 1: Figure S2). To assemble a collection with genomic and AST data, we retrieved samples from the NCBI Pathogen Detection AST Browser and Bacterial and Viral Bioinformatics Resource Center (BV-BRC) (Additional file 2: Table S5). First we deduplicated genomes from both sources using the BioSample accession, then assembly quality information was subtracted from the AllTheBacteria v2 [13] database and we kept only samples with a unique run accession and classified as *E. coli* by Sylph [34]. Samples falling outside ±2 standard deviations from the mean for *E. coli* taxonomic abundance (Sylph), total genome length, or contig count were discarded. We retrieved raw reads with fasterq-dump from the sra-toolkit v3.2.1 [12], and followed the same methodology as for *E. faecium* genomes to test the phenotype-genotype determination using: ARIBA v2.14.6, and AMRFinderPlus v4.0.23 [31], and SRST2 v0.2.0 [28], and SRST2-dev. We also tested SRST2-dev with an extended database. Guidelines and MIC cutoffs used to interpret AST are detailed in Table S2, Additional file 2. As there were no The European Committee on Antimicrobial Susceptibility Testing (EUCAST), Clinical & Laboratory Standards Institute (CLSI) or epidemiological cut-off values (ECOFFs) breakpoints for streptomycin in *E. coli*, we used the MIC distribution along with the presence of known streptomycin resistance genes to set the cutoff that clearly split the wildtype population from those with higher MICs and streptomycin resistance genes (i.e. MIC >32) (Additional file 1: Figure S3).

## Results

### E. faecium strains with duplicated and interrupted tet(M) result in tetracycline genotype-phenotype discrepancies

To identify the causes of tetracycline genotype–phenotype discrepancies in *E. faecium*, an antibiotic for which we had previously observed particularly low accuracy in genotypic resistance detection [3], we analyzed 2,610 *E. faecium* short-read genomes with curated tetracycline susceptibility profiles and AMRFinderPlus to detect tetracycline ARGs. Across this dataset, *tet*(M) was the most prevalent tetracycline resistance gene detected in 1,955 genomes, followed by *tet*(L) (n = 1,371) and *tet(S)* (n = 333). Co□occurrence of multiple *tet* genes was common, with 1,258 (48.1%) isolates carrying both *tet*(M) and *tet*(L) (Figure 2A).

**Figure 2.**
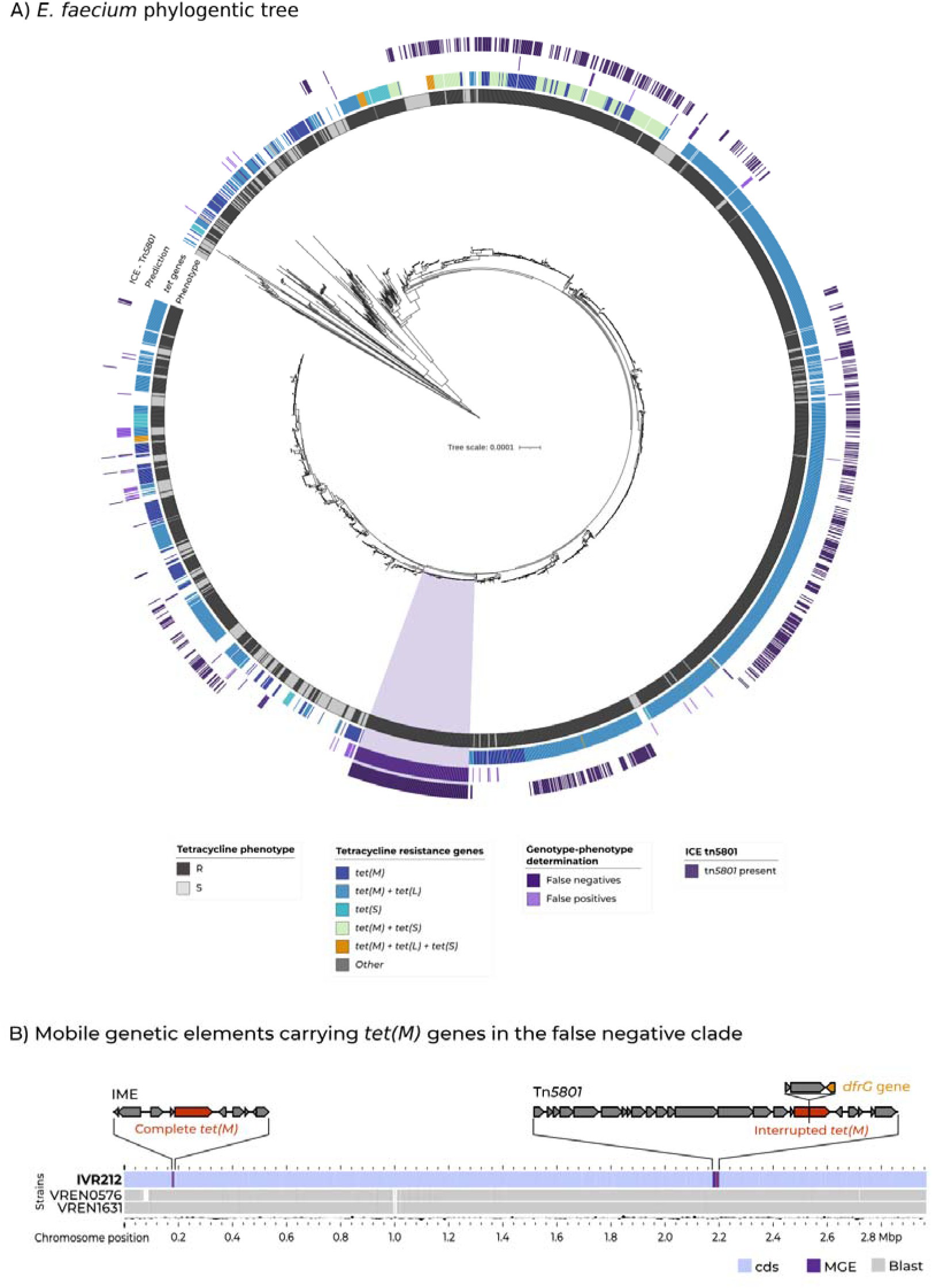
Phylogenetic context of tetracycline genotype–phenotype discrepancies in E. faecium. (A) Maximum□likelihood phylogeny of the 2,610 E. faecium Clade A isolates with tetracycline susceptibility data. The first ring indicates the AST phenotype (resistant or susceptible). The second ring shows the presence of tet(M), tet(L), tet(S), and other tet genes as detected by AMRFinderPlus using a stringent 98% coverage threshold. The third ring displays the genotype□based predictions for tetracycline resistance, highlighting false□positive and false□negative calls. The outer ring marks the presence of the Tn5801□like integrative and conjugative element (ICE) associated with interrupted tet(M) genes. (B) Mobile genetic elements carrying tet(M) in isolates from the false□negative clade. Shown is the genomic context of strain IVR212, which carries two tet(M) copies: a complete tet(M) gene located on an integrative mobile element (IME), and an interrupted tet(M) gene located on a Tn5801□like ICE, disrupted by a dfrG cassette. Genome comparisons with two additional long□read–sequenced isolates (VREN0576 and VREN1631) from the same false□negative lineage. Blast results show that both MGEs and the interrupted tet(M) configuration are conserved across the clade.

After excluding rare allelic variants associated with susceptibility and using a 98% gene coverage threshold for *tet* gene detection that maximized both sensitivity and specificity for detection of tetracycline resistance (Additional file 1: Supplementary Results and Figure S4), we assessed the accuracy of AMRFinderPlus to predict phenotype tetracycline resistance. Specificity was 89.4% and sensitivity 91.9%, with 178 false-negative isolates identified, that is, phenotypically resistant isolates lacking complete *tet* genes in their genomes. Notably, 138 out of 178 (77.5%) of these false negatives formed a wellOsupported monophyletic clade of resistant isolates that consistently carried a partially detected *tet*(M) gene (41.78% coverage, Figure 2A).

To investigate this clade, we selected three representative strains for ONT long-read sequencing (Figure 2B) and generated hybrid assemblies with their Illumina short-read data. All three hybrid assemblies contained a circularized chromosome of 2.9 Mb and 4-8 plasmids. All three genomes contained two copies of *tet*(M): a complete copy (100% coverage) located on an Integrative Mobile Element (IME), and an interrupted copy within a Tn*5801*-like Integrative and Conjugative Element (ICE) where *tet*(M) was split by a *dfrG* cassette into two partial fragments of 41.78% and 58.69% of coverage (Figure 2B). Both IME and ICE were integrated into the chromosome and present in all isolates belonging to the false-negative clade, while the ICE was also present in 60% of the entire genome collection (Additional file 1: Supplementary methods and supplementary results).

Given these observations, we concluded that the simultaneous presence of both an intact and fragmented tet(M) copies affected the detection of this gene from short□read sequence data alone.

### Duplicated and partial ARGs are common in E. faecium genomes across antibiotic classes

To determine whether this observed effect was restricted to tetracyclines or not, we screened a dataset of 823 high-quality closed *E. faecium* genomes (Additional file 1: Table S3) for ARG duplication and completeness across antibiotic classes. Among all genes, *tet*(M) was the most frequently duplicated or incomplete gene (n = 326 genomes), with 56.7% of isolates carrying *tet*(M) showing duplicated or incomplete copies. This pattern was primarily driven by the high prevalence of the Tn*5801*□like ICE carrying the interrupted *tet*(M) copy in the collection (Additional file 1: Supplementary results and figure S5). We also detected other resistance genes frequently duplicated or incomplete, including *erm*(B) (responsible for macrolide resistance), and *ant(6)-Ia* and *aph(3’)-IIIa* (aminoglycoside resistance) (Figure 3 A). These genes were predominantly found duplicated with 84.8% (n=123/145), 75.4% (n=83/110), and 94.1% (n=48/51) of genomes carrying two or more complete copies, respectively.

**Figure 3.**
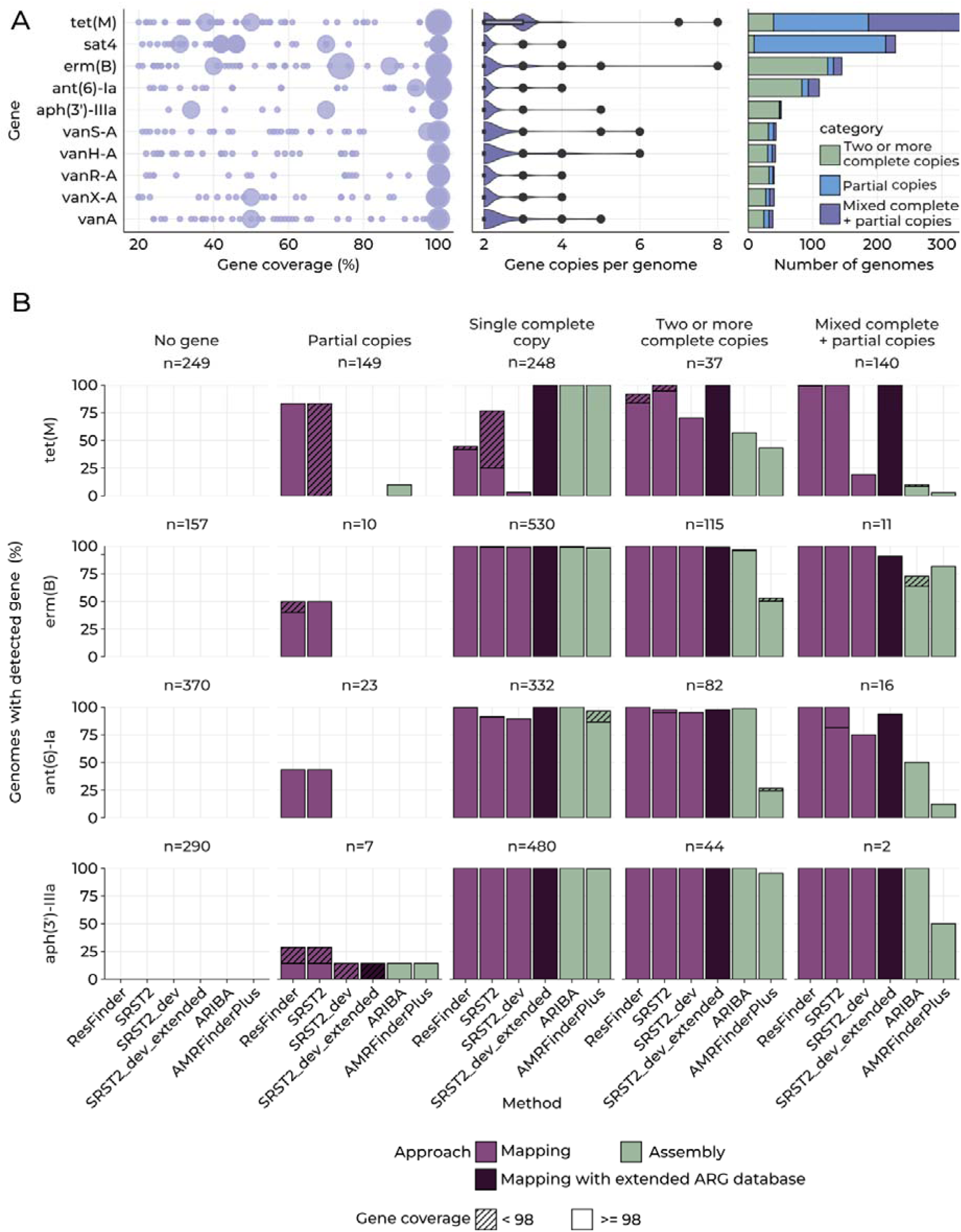
Overview of duplicated and partial ARGs in E. faecium closed genomes and performance of detection tools across structural configurations. (A) Identification of duplicated and partial ARGs in 823 closed E. faecium genomes. For the ARGs most frequently present in multiple copies, the distribution of coverage values across genomes is shown. Violin plots display the number of gene copies per genome, together with the proportion of genomes containing two or more complete copies, mixed configurations of partial and complete copies, or only partial copies for each ARG. (B) Evaluation of ARG detection across structural configurations using short□read data. ARG structural categories defined from the closed genomes were compared to detection outcomes obtained from IlluminaOsimulated reads. Two mappingObased tools (ResFinder, SRST2), two assemblyObased tools (ARIBA, AMRFinderPlus), and a modified version of SRST2 were tested, using either the standard AMRFinderPlus database or an extended database including additional allelic variants. For each method (xOaxis) and each structural category, stacked bars show the proportion of genomes in which the tool reported the ARG.

### Variables affecting the bioinformatic detection of ARGs from Illumina sequence data

To evaluate how allelic diversity, completeness, and duplication of ARGs could affect their detection from Illumina short-read data, we simulated Illumina reads from closed *E. faecium* genomes and assessed the ability of commonly used ARG detection tools to call the ARGs identified in the closed genomes. (Figure 1 A). ARG calls in each closed genome were classified as: genomes containing only partial copies of the gene (coverage < 90%), genomes with a single complete copy (coverage over 90%), genomes with two or more complete copies, and genomes carrying both partial and complete copies of the same gene (Figure 3 B). We tested ResFinder, SRST2, ARIBA, and AMRFinderPlus (using Shovill generated assemblies) with default settings.

We first observed that AMRFinderPlus incorrectly identified 33 ARGs (1.4% of all AMRFinderPlus ARG calls) as partial despite being complete in the closed genomes. This affected all four ARGs analyzed, especially *ant(6)-Ia* (n = 16), *erm*(B) (n = 6), *tet*(M) (n = 5), and *aph(3’)-IIIa* (n = 3) (Additional file 1: Figure S6). These errors resulted from high sequence divergence between the samples’ ARG allelic variants and those represented in the AMRFinderPlus database. After manually curating these categorization errors, we observed a consistent pattern: the accuracy of ARG identification was mostly influenced by the detection approach (mapping vs. assembly) and the structural configuration of the gene. Mapping-based tools (i.e., ResFinder and SRST2) frequently misclassified partial genes as complete, ranging from 23% to 83.2% of misclassification depending on the ARG (see Figure 3B, “Partial copies”). For *tet*(M), mapping based tools misclassified 124 of 149 genomes with partial genes as complete. These misclassifications were due to these tools evaluating gene presence only based on the overall mapping coverage of short reads and not assessing patterns in the short-read alignments that indicate interrupted genes (such as clipped reads or coverage drops), as it is the case for the interrupted *tet*(M) in the ICE (Figure 2B).

Additionally, mapping-based tools underestimated ARG presence in genomes with a single complete copy, particularly for *tet*(M) and *ant(6)-Ia* (Figure 3B, “Single complete copy”). These failures were largely due to insufficient allele representation in the AMRFinderPlus database. Only 4 of the 18 *ant(6)-Ia* alleles found in *E. faecium* closed genomes were included in the AMRFiderPlus database, and only 42 of 75 for *tet*(M). As a result, reads from divergent alleles mapped poorly and failed to meet coverage thresholds required for reporting complete genes.

Assembly-based tools (i.e. ARIBA and AMRFinderPlus) showed a distinct error pattern, failing to detect complete copies in genomes with duplicated or mixed copies of partial and complete ARGs. Across ARGs, AMRFinderPlus detected only 51% of duplicated copies overall (range 27.5–95.5%) and 9.8% mixed configurations (range 3.0 - 81.8%). ARIBA performed well for duplicated ARGs (overall 93.1% correctly detected, range 56.8 - 100%) but dropped sharply in the detections of mixed ARG configurations (19.5% overall, range 10.4 - 100%). These patterns indicate that the presence of multiple copies of ARGs, whether complete or partial, generate repetitive regions that cannot be resolved by short-read assemblers, preventing the accurate assembly and detection of these genes.

### Bioinformatic improvements in the detection of ARGs from Illumina sequence data

Having flagged the limitations of short-read assembly methods to detect certain configurations of ARGs, we aimed to improve short-read mapping detection methods. Specifically, we modified SRST2 [28] (SRST2_dev, modification available in https://github.com/NerisGarcia/srst2_NGG) to identify and flag potential gene interruptions by detecting abrupt coverage drops or the accumulation of clipped reads within the ARG short-read alignment (Additional file 1: Figure S7). We also increased the allele representation in the AMRFinderPlus ARG database including 33 new *tet*(M) alleles (total 75 variants), 10/20 additional *tet*(L), 5/7 *aph(3’)-IIIa,* 14/18 *ant(6)-Ia* and 21/35 *erm*(B) variants identified in the *E. faecium* closed genomes. No new alleles were identified for *tet*(S). To assess the impact of this extended database, we also compared the performance of our modified SRST2 pipeline (here called SRST2-dev) with and without using it.

In comparison to existing tools, SRST2-dev with the extended database showed a marked increase in detection of complete copies when duplicated or in mixed configurations, reporting 98.9% and 98.8% of cases, respectively (Figure 3 B). We also found that increasing ARG allelic representation had a major impact on detection accuracy for specific ARGs, more notably *tet*(M) and *ant(6)-Ia*. For example, the same SRST2-dev pipeline without the extended database failed to detect *tet*(M) in nearly all categories, reporting only 8 single-copy ARGs compared to 248 with the extended database. Our stricter evaluation of read alignment in SRST2-dev rejects distant alleles due to poor mapping quality, thus requiring better allelic representation in the ARG database to detect all ARGs.

### Impact of gene duplication and completeness on genotype-phenotype determination across antibiotic classes

We next assessed how allelic diversity, completeness, and duplication of ARGs could affect the accuracy of phenotypic resistance prediction in real *E. faecium* clinical strains. To do this, we used a curated short□read dataset of 4,507 *E. faecium* genomes with corresponding AST results [3] (Additional file 2: Table S1). We focused on antibiotics associated with duplicated and partial ARGs that had both sufficient sample size (n=2,000) and a high prevalence of genomes carrying the studied gene as the only resistance marker (Figure 1 B). Antibiotics included were tetracycline (*tet*(M), 2,438 isolates tested), erythromycin (*erm*(B), 2,250 isolates), kanamycin (*aph(30)-IIIa,* 2,356 isolates), and streptomycin (*ant(6)-Ia,* 2,353 isolates).

Accounting for gene duplication and completeness increased sensitivity across multiple classes but introduced specificity losses majorly related to specific gene variants or lineages (Figure 4). For tetracycline, where truncation and duplication of *tet*(M) affects detection, SRST2-dev with the extended database achieved the best overall accuracy (97.7%, 95% CI 97.03–98.26%) and highest sensitivity (99.75%, 95% CI 99.42–99.92%). Low specificity (87.97%, 95% CI 84.49–90.91%) was due to 51 false positives (other tools ranged from 38 to 165). These false positives were strongly enriched for specific ARG alleles: 37.2% (n=19) carried a *tet*(L)*08* allele associated to reduced susceptibility (Additional file 1: Figure S4) whereas 41.1% (n=21) corresponded to *tet*(M) alleles with aminoOacid deletions. This suggests that the mismatch likely reflects nonOfunctional or partially functional variants, rather than detection errors. We also observed differences in specific *tet* genes reported by each tool (Figure 4). SRST2OdevOextended identified most isolates as carrying *tet*(M) (with or without *tet*(L)), whereas AMRFinderPlus and ARIBA frequently reported only *tet*(L) in these isolates due to the lack of detection of duplicated *tet*(M).

**Figure 4.**
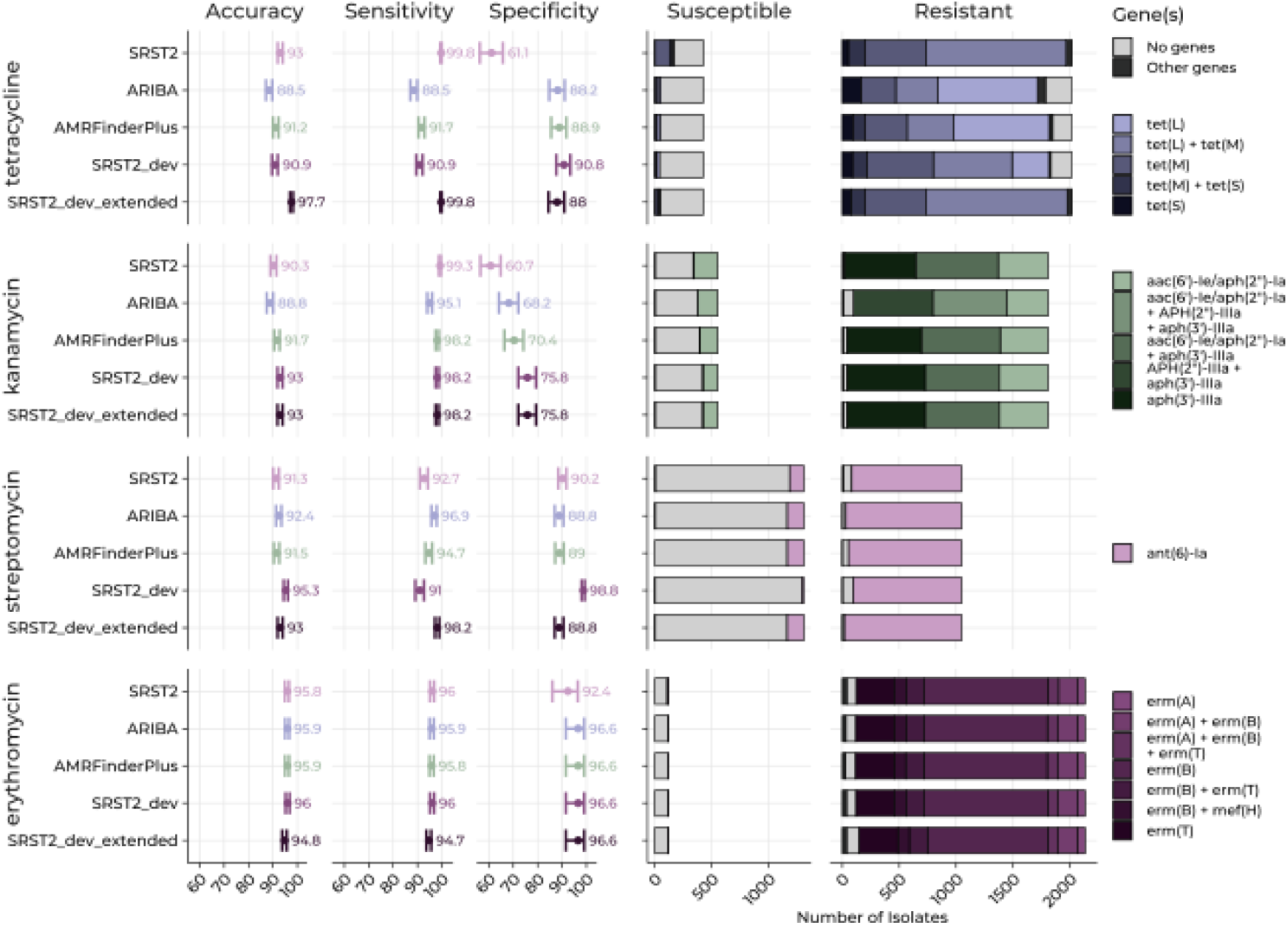
Phenotype determination from ARG detection and ARG reporting across tools in E. faecium. The left panel shows the sensitivity, specificity, and diagnostic accuracy for each antibiotic–tool combination in the E. faecium clinical dataset. Each point represents the estimated value of the metric, and the whiskers correspond to the 95% confidence intervals. The right panel shows the distribution of reported ARGs across tools and methods among phenotypically susceptible and resistant E. faecium isolates.

Detection of kanamycin resistance (attributable in part to *aph(3’)-IIIa*) also benefited from the SRST2-dev extended pipeline, which yielded the highest specificity among all tested tools (75.82%, 95% CI 72.02-79.34%). Despite this improvement, 133 false positives remained, also shared across all tools (range 133–216). Most of these (128, 96.2%) also harbored the *aac(6’)0Ie/aph(2’’)0Ia* gene and clustered phylogenetically (Additional file 1: Figure S8), which suggests the existence of lineage-specific variants associated with susceptibility instead of AST or ARG detection errors.

In contrast, for streptomycin resistance detection (attributable in part to *ant(6)-Ia*), SRST2-dev tool outperformed the version using the extended database (95.33% vs. 92.99% accuracy). Using the extended database increased sensitivity (90.9% to 98.1%) at the expense of specificity (98.78% to 88.83%). The additional FPs (n=130) detected by SRST2-dev with the extended database were due to detection of new *ant(6)-Ia* variants, a significant portion of these (42.3%, n=55) grouped within a single clade (Additional file 1: Figure S9), again indicating ARG alleles associated with susceptibility rather than bioinformatic failures. The other false positives and false negatives were mostly grouped in the A2 clade.

Finally, for erythromycin resistance detection (associated mostly to erm(B)), we found comparable high performance across tools (Figure 4), indicating that structural variation and allelic diversity in erm(B) have minimum impact on genotypic detection of erythromycin resistance.

### ARG duplication also affects AMR determination from Escherichia coli genomes

To test whether these effects were species□specific or more generalizable, we applied the same methodology to *E. coli.* Using 4,774 complete *E. coli* genomes (Additional file 2: Table S4), we identified duplicated and partial ARGs affecting resistance to tetracyclines (*tet*(A)), sulfonamides (*sul1, sul2*), streptomycin (*aph(6), aph(3)*), chloramphenicol (*floR*), and aminoglycosides (*aadA2*) (Figure 5 A). We then assessed whether duplication, completeness and allelic diversity of ARGs influenced genotype–phenotype prediction in a curated short□read dataset of 7,437 *E. coli* genomes with available AST data (Additional file 2: Table S5). Analyses focused on tetracycline (*tet(A)*, 5,863 isolates), streptomycin (*aph(6)-Ia/aph(3’’)-Ib*, 2,842 isolates), and chloramphenicol (*floR*, 5,284 isolates) as these antibiotics had both sufficient sample size (n=2,000) and a high prevalence of genomes carrying the studied gene as the only resistance marker.

**Figure 5.**
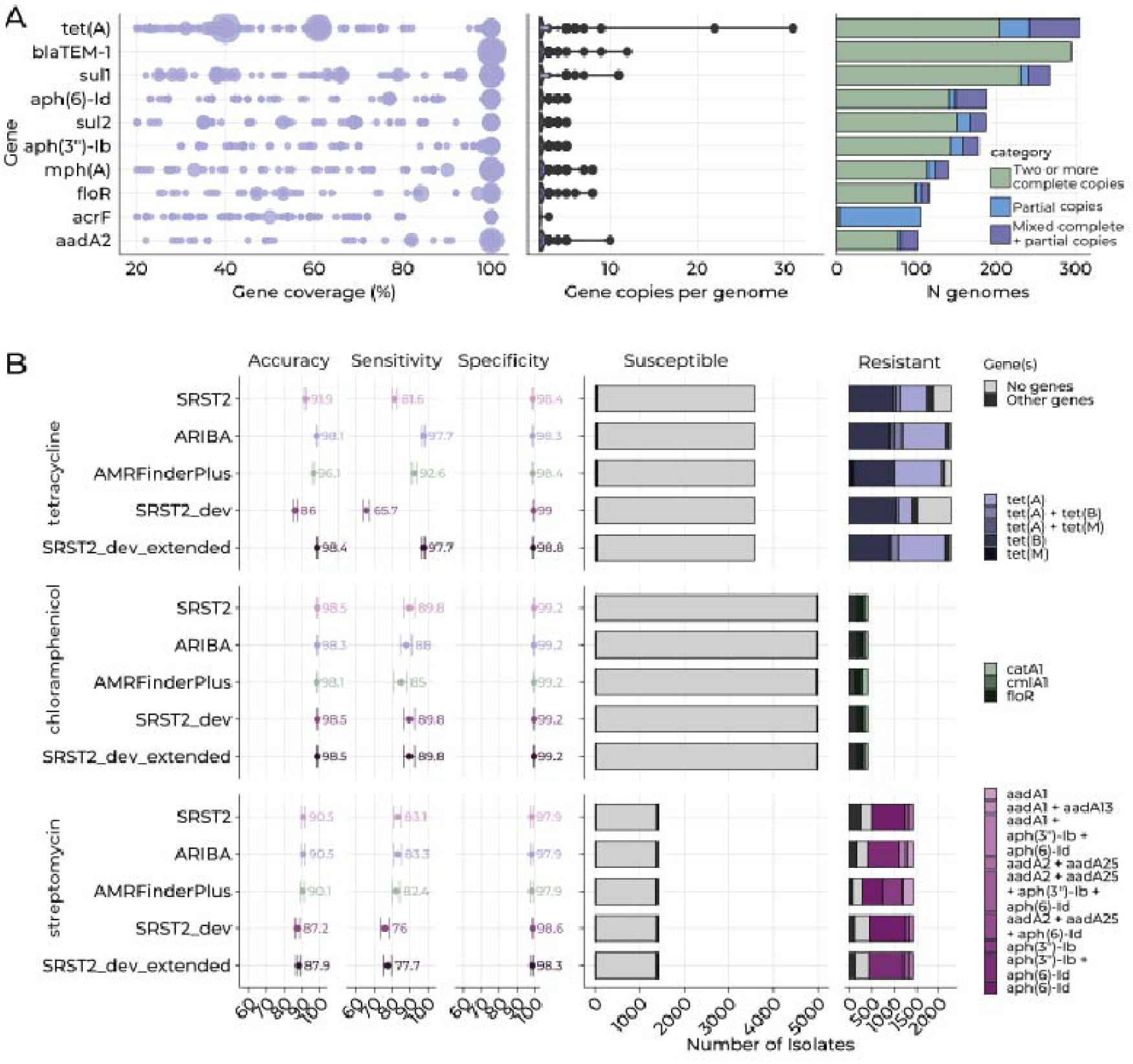
Detection of duplicated and partial ARGs and phenotype determination from ARG detection in *E. coli*. (A) Detection of duplicated and partial ARGs in 4,774 *E. coli* closed genomes. The ARGs most frequently found in multiple copies are shown. For each ARG, the distribution of coverage values across genomes is displayed, together with violin plots showing the number of ARG copies per genome and the proportion of genomes containing two or more complete copies, mixed configurations of partial and complete copies, or only partial copies. (B) Accuracy metrics and ARG reporting in the *E. coli* clinical dataset. The left side shows the sensitivity, specificity, and diagnostic accuracy obtained for each antibiotic–tool combination. Each point represents the estimated value of the metric, and the whiskers indicate the corresponding 95% confidence intervals. The right side displays stacked bars showing the number of isolates in which each tool reported the relevant ARG, stratified by phenotypically susceptible and resistant isolates.

In *E. coli*, the effect of ARG variation on AMR genotype–phenotype concordance was most evident for tetracycline. The use of SRST2-dev with the extended database of ARG alleles led to the largest improvement in accuracy to this antibiotic (Figure 5B), underscoring the importance of allele representation in ARG databases. For chloramphenicol, predictions were highly consistent across methods. The low number of chloramphenicol resistant isolates (n=401) might explain in part the apparent lack of influence of the factors assessed here (ARG duplication and allelic diversity). For streptomycin, all tools displayed lower sensitivity than for other antibiotic classes. AMRFinderPlus, ARIBA, and SRST2 performed similarly (82–83% sensitivity; ∼90% accuracy), but SRST2-dev with the extended database show lower values (87.9% of accuracy) suggesting only a modest impact of allelic diversity on *ant(6)0Ia* detection in *E. coli* compared to its strong effect in *E. faecium*.

## Discussion

Here we showed that completeness, copy number, and allelic diversity of ARGs can impact the accurate detection of ARGs from short-read data and influence genotype–phenotype agreement for certain antibiotics. Although originally investigated here for tetracycline resistance in *E. faecium*, we showed that these factors affect multiple antibiotic classes and other organisms like *E. coli*. We also implemented a bioinformatics strategy to account for ARG allelic diversity and handle the detection of duplicated and incomplete ARGs from Illumina short-read data.

Previous work proposed allelic variation [7] and silenced acquired ARGs [8] as potential drivers of low specificity in genotypic AMR detection (phenotypically susceptible isolates carrying ARGs). Thus, we started testing the association of individual *tet* alleles with phenotypic susceptibility. Despite a single *tet*(L) variant significantly enriched among susceptible isolates, allelic diversity did not explain the discrepancies observed. In contrast, we found that the coverage thresholds used for ARG gene detection had a strong impact on genotype-phenotype agreement. Adjusting to an empirical coverage threshold for *tet* genes substantially improved specificity, showing that generic default thresholds may be inadequate, and that ARG- and organism-specific thresholds should be established and used instead.

After accounting for allelic variation and gene coverage, sensitivity in predicting tetracycline resistance was still low, mostly attributable to a single clade of false negatives (phenotypically tetracyclineOresistant isolates lacking a complete *tet*(M) detected). Long-read sequencing revealed that isolates in this clade carried both a complete copy of *tet*(M) on an IME and an interrupted copy on an ICE. We found that the coexistence of both a complete and partial copies of *tet*(M) led to partial recovery or loss of the complete gene from short-read assemblies, pointing to a bioinformatic artifact rather than a genetic basis for this AMR genotype-phenotype discrepancy.

Having identified that gene completeness and duplication may be a key driver of genotype-phenotype discrepancies for tetracyclines, we next investigated whether this was a broader feature of the *E. faecium* resistome. Analysis of closed genomes showed that multiple ARGs are frequently duplicated or present in mixed configurations of complete and partial copies. Next, we simulated Illumina short reads from closed genomes to evaluate the performance of different bioinformatic tools in detecting different ARG configurations with special focus on detection of gene interruptions and duplications, and how this may affect AMR phenotype determinations. We showed that current AMR detection approaches struggle with certain ARGs configurations in different ways. Mapping-based tools like ResFinder and SRST2 may classify interrupted genes as complete, thus overcalling resistance, because they rely on global gene coverage metrics and do not assess alignment artifacts such as soft-clipped reads or local coverage drops. In contrast, AMR tools based on short-read assemblies, such as ARIBA and AMRFinderPlus, frequently fail to recover complete genes when multiple or partial copies are present in the genome, thus undercalling resistance. The latter configuration creates repetitive regions that short-read assemblers cannot resolve, leading to contig collapse or fragmentation, which prevents the accurate reconstruction of the full ARG [35]. While duplication has been previously linked to AMR detection errors in *E. coli* [9,36], our results demonstrate that it is both common and directly impacts AMR genotype– phenotype agreement.

Finally, we also explored how allele representativeness in ARG databases could affect genotype–phenotype discrepancies. As previously reported [37], we found that including representative alleles is key. Even in closed genomes, AMRFinderPlus misclassified complete genes as partial due to divergence between alleles in strains’ genomes and those in the database. In general, extending the database with species-specific alleles improved detection across tools, particularly for mapping-based approaches [9]. Still, expanding allele representativeness in AMR databases alone is not sufficient unless allele-level interpretation of resistance is implemented to correctly link genotype to phenotype [7,9,31,37–39].

By extending our analysis to *E. coli,* we showed that these challenges are not specific to *E. faecium*. In *E. coli*, ARGs related to tetracyclines, sulfonamides, aminoglycosides, and chloramphenicol were frequently duplicated or incomplete in closed genomes, in some instances influencing AMR genotype–phenotype agreement. Our findings are consistent with prior evidence showing that duplicated ARGs are widespread among bacteria, especially in strains of clinical and livestock origin [40]. Altogether, these observations indicate that ARG detection errors arising from duplicated or structurally complex genomic contexts are likely to be common in bacteria if ARG integrity, copy number, and allelic variation are not accounted for.

Future tools and updates should account for these limitations. First, ARG gene and allele databases should be systematically expanded to capture the allelic diversity observed across species, leveraging variation already present in publicly available genomes. Second, tools should incorporate flags indicating potential gene duplication or truncation. Mapping-based approaches could compare ARG read depth to that of core chromosomal genes to identify increases in coverage depth consistent with duplications, and use read clipping patterns to detect interruptions or truncations. Assembly-based methods could be complemented with read-mapping steps to validate gene calls and assess the same signals from read alignments. Ultimately, future tools should be reporting ARG integrity, copy number, and their corresponding alleles.

This study has several limitations. Although our bioinformatics solution (modified mapping-based SRST2) improved detection of the challenging ARG configurations examined here, further development and validation may be required for other ARG families and organisms. In real genomic datasets, as the ones used in this study, multiple acquired ARGs frequently coOoccur, making it difficult to isolate the impact of improved detection of individual ARGs on sensitivity and specificity, and some collections may not contain duplicated or interrupted genes for certain antibiotics, limiting the ability to observe these effects. Our analysis focused solely on the detection of acquired ARGs and did not incorporate allelic variants associated with phenotypic susceptibility, which can confound interpretation. Finally, for the clinical isolates used in phenotype determination, we did not have both short-read and long-read sequence data, which would have enabled precise identification of the causes of genotype–phenotype discrepancies. Future benchmarking efforts should therefore employ datasets with both shortO and longOread sequence data.

## Conclusions

In conclusion, our results show that reliable genomic determination of AMR requires moving beyond detection of single-copy ARGs and alleles toward evaluation of AMR gene copy number, integrity, and allelic variation in a species- and antibiotic-specific manner. Incorporating these factors into genotypic AMR detection pipelines would improve genotype–phenotype agreement, and improve the accuracy and reliability of genomic surveillance strategies.

## List of abbreviations

AMR: Antimicrobial Resistance
AST: Antimicrobial Susceptibility Testing
WGS: Whole Genome Sequencing
CLSI: Clinical & Laboratory Standards Institute
MIC: Minimum Inhibitory Concentration
SRA: Sequence Read Archive
NCBI: National Center for Biotechnology Information
BV-BRC: Bacterial and Viral Bioinformatics Resource Center
WSI: Wellcome Sanger Institute
EUCAST: The European Committee on Antimicrobial Susceptibility Testing
ECOFFs: Epidemiological cut-off values
IME: Integrative Mobile Element
ICE: Integrative and Conjugative Element

## Declarations

### Ethics approval and consent to participate

Not applicable.

### Consent for publication

Not applicable.

### Availability of data and materials

Genome sequence data analyzed in this study were obtained from publicly available repositories, including the NCBI databases and the Sequence Read Archive (SRA). Accession numbers for all publicly available Illumina short-read datasets and genome assemblies used in the analyses are provided in Additional file 2 (Supplementary Tables). Oxford Nanopore Technologies (ONT) genome sequences generated for this study have been deposited in the NCBI Sequence Read Archive under BioProject PRJNA1456658. Corresponding BioSample accession numbers are as follows: VREN0576 (SAMN57420212), VREN1631 (SAMN57420326), and IVR212 (SAMN57443460). Antimicrobial susceptibility phenotype data are publicly available from the original sources and are also compiled in the supplementary tables associated with this article. All custom code developed for this study is publicly available. Analysis scripts are available at https://github.com/NerisGarcia/Tet_resistance, and the modified version of the SRST2 software used in this work is available at https://github.com/NerisGarcia/srst2_NGG.

## Authors’ contributions

FC and NGG designed this study. NGG conducted all the bioinformatic analyses with contributions from FC, RF and JIK. RF curated the dataset of public *E. coli* genomes with available phenotypic AST data. BB conducted the DNA extractions needed to perform ONT sequencing. JIK performed the bioinformatic analyses related to the generation of Illumina-ONT hybrid assemblies. MET, EMH and TG contributed to resources and funding for the ONT sequencing. FC and NGG wrote the first version of the manuscript with contributions from all other authors. NGG created all the figures and tables in this manuscript. All authors read and approved the final version of the manuscript.

## Acknowledgements

The authors thankfully acknowledge RES resources provided by University of Málaga in Picasso HPC (activities BCV-2024-3-0018 and BCV-2025-1-0012).

## Additional files

*Additional file 1.* Supplementary methods, results, and figures.

*Additional file 2.* Supplementary tables.

Table S1: *Enterococcus faecium* real data collection. Accessions, clade, AST;

Table S2: Guidelines and MIC cutoff used to interpret AST data

Table S3: Enterococcus faecium closed genome collection;

Table S4: Escherichia coli closed genome collection;

Table S5: Escherichia coli real data collection.

